# Early identification of dengue virus lineage replacement in Brazil using portable genomic surveillance

**DOI:** 10.1101/716159

**Authors:** Jaqueline Goes de Jesus, Karina Rocha Dutra, Flavia Cristina da Silva Salles, Ingra Morales Claro, Ana Carolina Terzian, Darlan da Silva Candido, Sarah C. Hill, Julien Thézé, Tatiana Lang D’Agostini, Alvina Clara Felix, Andreia F. Negri Reis, Luiz Carlos Junior Alcantara, André L. Abreu, Júlio H. R. Croda, Wanderson K. de Oliveira, Ana Maria Bispo de Filipis, Maria do Carmo Rodrigues dos Santos Camis, Camila Malta Romano, Nick J. Loman, Oliver G. Pybus, Ester Cerdeira Sabino, Mauricio L. Nogueira, Nuno Rodrigues Faria

## Abstract

Over 400 million people are estimated to be at risk of acquiring dengue virus (DENV). Despite efforts to mitigate the impact of DENV epidemics, the virus remains a public health problem in the Americas: more than one million DENV cases were reported in the continent between January and July 2019 DENV was first detected in Brazil in 1982, and Brazil has reported 88% (1,127,244 cases) of all DENV cases in the Americas during 2019 to date. São Paulo state in the southeast of Brazil has reported nearly half of all DENV infections in the country. Here we characterised the genetic diversity of DENV strains circulating in São Paulo state in 2019, at the epicentre of the ongoing DENV epidemic. Using portable nanopore sequencing we generated 20 new DENV genome sequences from viremic patients with suspected dengue infection residing in two of the most-affected municipalities, Araraquara and São José do Rio Preto. We conducted a comprehensive phylogenetic analysis with 1,630 global DENV strains to better understand the evolutionary history of the DENV lineages that currently circulate in the region. The new outbreak strains were classified as DENV2 genotype III (American/Asian genotype). Notably, phylogenetic analysis indicated that the 2019 outbreak is the result of a novel DENV lineage that was recently introduced to Brazil from the Caribbean region. Our genetic analysis further indicates that the introduction and onwards spread of the outbreak lineage (named here DENV2-III BR-4) indicates a new DENV2 lineage replacement in Brazil. Dating phylogeographic analysis suggests that DENV2-III BR-4 was introduced to Brazil in or around early 2014, possibly from the Caribbean region. Our study describes the early detection of a newly introduced and rapidly-expanding DENV2 virus lineage in Brazil.

**Author Summary:** Dengue is the most important mosquito-borne viral disease of humans. The disease is caused by the dengue virus (DENV) that is classified within genus *Flavivirus*. DENV infections are caused by 4 serotypes (DENV 1-4) that are genetically related but antigenically distinct. Dengue infection results in a variety of symptoms that range from mild fever to dengue hemorrhagic fever and/or dengue shock syndrome (DHF/DSS). Clinical outcomes are associated with different types of infection, viral serotypes, genotypes, lineages, and host genetic factors. As a re-emerging infectious disease, DENV has become a serious threat to public health in the Americas, and particularly in Brazil, where it was introduced in the 1980s and became well established due to the country-wide re-infestation of the *Aedes aegypti* mosquito vector species. During the first six months of 2019, 1,282,183 DENV cases were reported in the Americas, with Brazil reporting a staggering 1,127,244 (88%) of all dengue cases in the continent. To date, no information exists on the genetic composition of the DENV lineage or lineages causing the current epidemic. Here we use portable sequencing to rapidly generate virus genome data from cases occurring in two different are severely-affected municipalities in São Paulo state, Brazil. We find that the 2019 dengue outbreak in Brazil is caused by a newly introduced DENV serotype 2 genotype III (Asian/American) that seems to be replacing previously-circulating DENV2 lineages. We discuss the potential implications of our results regarding the current outbreak in the context of previous outbreaks in the same region.

## Introduction

Over 400 million people are estimated to be at risk of acquiring dengue virus (DENV, genus *Flavivirus*, family *Flaviviridae*) [1], a mosquito-borne virus transmitted in tropical and subtropical areas by competent urban vectors such as the mosquitoes *Aedes aegypti* and *Aedes albopictus* [2]. DENV is classified into four distinct virus lineages named serotypes 1 to 4 (DENV1-4). Within each DENV serotype there is some degree of genetic variation, and at least 19 DENV genotypes have now been described [3]. Increasing human mobility has facilitated the co-circulation of multiple dengue serotypes in the same region [4], a pattern known as hyperendemicity. In such regions, DENV epidemiological dynamics are complex and typically characterized by virus genotype replacement every 7-10 years [5-9]. Clade replacement is typically associated with an increased number of cases and cases with severe disease. Certain genotypes and lineages seem to be more frequently associated with severe disease outcomes [8, 10, 11].

DENV was first detected in Brazil in 1982 [12]. Since then, it has become a serious public health concern due to its high incidence in the country and association with severe dengue illnesses [13]. Co-circulation of dengue serotypes has been observed throughout Brazil [14, 15], particularly in highly populated areas of the southeastern region that includes the federal states of São Paulo, Rio de Janeiro, Minas Gerais and Espírito Santo. Between 1995 and 2015, Brazil reported nearly 8 million DENV cases, which comprises 55% of all cases reported in the Americas during this period [12]. Over the last thirty years, the southeast region of Brazil has reported 2225 dengue-related fatal cases, representing 43% of all dengue-related deaths in the country [13].

In the first half of 2019 (1^st^ Jan to 30^th^ Jun) Brazil has already reported 1,127,244 dengue cases [12]. Importantly, this number is nearly 8-fold higher than in the previous year and corresponds to 89% of all the dengue cases reported in the Americas over the same period. The number of severe cases (n=710) and dengue-related deaths (n=366) also increased by at least 2.3-fold in comparison with 2018.

The Southeast region of Brazil has reported 65.7% of all dengue cases identified in the country [16]. São Paulo state is the most highly densely populated state and the main socio-economic hub in Brazil; previous studies suggest the state was an important source location for the spread of DENV4 in the country [17]. Here, we characterise the genetic diversity of circulating DENV in two municipalities of São Paulo state. We generated virus genome sequences from the ongoing outbreak using a well-established portable genomic approach [18, 19]. In an attempt to better understand the origin and dynamics of the 2019 outbreak, we conducted comprehensive genetic analyses to understand the relationship between the current epidemic strains and those that circulated in previous outbreaks in the Americas.

## Methods

Brazil is organized into 26 federal states and 1 federal district. Sao Paulo state is the most populous Brazilian state and comprises 615 municipalities. São José do Rio Preto is the 11^th^ most populated municipality (450,657 inhabitants), and Araraquara the 32^nd^ most populated municipality (230,770 inhabitants) in the state (www.ibge.gov.br). In each municipality, the number of dengue suspected cases is notified by local public health secretaries to the Centro de Vigilância Epidemiológica “Prof.Alexandre Vranjac” (CVE), part of Sao Paulo’s State Health Secretary. As part of dengue surveillance efforts in São Paulo state, samples are collected from patients suspected of acute DENV infection and tested for DENV by real-time quantitative reverse transcription PCR (qRT-PCR) by several research centers and public health institutions, including Adolfo Lutz Institute. Monthly numbers of dengue cases per serotype are then aggregated by CVE.

To assess the genetic diversity of dengue cases circulating in Sao Paulo state, we selected 20 qRT-PCR positive samples DENV serotype 2 from patients in two municipalities, Araraquara and São José do Rio Preto. The majority of the samples (19 out of 20) was collected between January and the end of April 2019; however, to investigate whether the same lineage was circulating before 2019, we also included one sample from São José do Rio Preto collected in early June 2017. Samples had mean RT-qPCR cycle-threshold values of 19.8 (range: 16.4–25). Diagnostic details and symptoms are shown in **Table S1**.

Residual anonymized clinical diagnostic samples from Araraquara were obtained following ethical approval by Hospital das Clínicas – University of São Paulo’s Institutional Review Board (CAPPesq) (number 3.156.894). São José do Rio Preto samples were obtained from virological surveillance routine, within the study approved by University of São José do Rio Preto Institutional Review Board approval #48982/2012. We used residual anonymized clinical diagnostic samples, with no risk to patients, which were provided for research and surveillance purposes within the terms of Resolution 510/2016 of CONEP (Comissão Nacional de Ética em Pesquisa, Ministério da Saúde; National Ethical Committee for Research, Ministry of Health).

The 20 qRT-PCR–positive DENV2 samples were subjected to viral genomic amplification at the Institute of Tropical Medicine, University of São Paulo, Brazil. Genome sequencing was conducted using the portable nanopore MinION sequencing platform, which has been used previously in Brazil during outbreaks of Zika virus and yellow fever virus [9-11]. Sequencing was performed using a multiplex PCR primer scheme designed to amplify the entire coding region of DENV2 as previously described [12].

RNA was extracted and reverse-transcribed to cDNA using Superscript IV First-Strand Synthesis System (Thermo Fisher Scientific, MA, US) and random hexamer priming. Then, multiplex PCR was performed to generate overlapping amplicons of the whole genome of the targeted DENV2 strain. DENV2 genome amplification consisted of 35 cycles of PCR according to the reaction mix and thermocycling described by Quick et al [19]. AmpureXP purification beads (Beckman Coulter, High Wycombe, UK) were used to clean up PCR products, which were then quantified by Qubit dsDNA High Sensitivity assay on a Qubit 3.0 instrument (Life Technologies). Sequencing libraries were generated using the Genomic DNA Sequencing Kit SQK-LSK108 (Oxford Nanopore Technologies), by pooling, in equimolar proportions, a total of 250 ng of PCR products previously barcoded using the Native Barcoding Kit (NBD103, Oxford Nanopore Technologies, Oxford, UK). The libraries were loaded onto an Oxford Nanopore flow cell R9.4 (FLO-MIN106) and sequencing data were collected for 30 hours. The median number of mapped reads was 45,013 reads per sample, and the generated consensus genomes had a mean coverage of 81% of the genome at 20x minimum sequencing depth. Sequencing statistics for each sample are shown in **Table S1**. Raw and processed data are available on GitHub (https://github.com/arbospread).

To investigate the origins of the newly generated genomes we downloaded all DENV2 nucleotide sequences longer than 1400 nucelotides (nt) from GenBank [16] that had a known date (year, and month and day when available) and location (country, and city when available; n=1630 as of 20 June 2019). We aligned these sequences using MAFFT automatic settings [18] and manually edited them with AliView v1.19 [19]. We subsequently constructed an initial maximum likelihood phylogeny to help identify the genotypes of strains that have historically circulated (collected before 2019) and are currently circulating (collected in 2019) in Brazil. For this genotype assessment, we constructed phylogenies using FastTree v.2 with gamma-distributed among site rate heterogeneity and a general time reversible nucleotide substitution model [20]. We observed that all sequences from the Americas (including the newly generated sequences) grouped together in a well-supported monophyletic clade. Therefore we next constructed a dataset comprising only sequences collected in the Americas (n=670). To reduce sampling bias towards a high number of samples from well-sampled countries, we removed duplicate sequences (same day and location) from Nicaragua and Peru, yielding a final dataset of 436 genomes (including 66 genomes collected in Brazil between 1990 to 2013). Maximum likelihood phylogenies of the American DENV2 genomes (n=670 and n=436) were generated using PhyML [20] available through Seaview v.4.6.1, using gamma-distributed among site rate heterogeneity and a general time reversible nucleotide substitution model [21]. Root-to-tip divergence and temporal signal was evaluated using TempEst [21].

Georefenced and time-stamped phylogenies were constructed using a discrete phylogeographic approach as previously described [19, 22]. In brief, countries were grouped into four geographic regions consisting of Brazil (n=86), Central America and Mexico (n=45), South America (n=132) and Caribbean (n=173). Inferred locations at each internal node and corresponding dated phylogenies trees were estimated using BEAST1.10 [23]. MCMC convergence was inspected using Tracer.v1.7 and summary trees were generated using TreeAnnotator [23].

## Results and Discussion

Epidemiological information on the number of confirmed dengue cases with associated serotype information shows that all four dengue serotypes co-circulate in Sao Paulo state (**Figure 1A**). While in 2012 DENV1 (52%, n=373/712) and DENV4 (39%, n=274/712) predominated, DENV 2 (9%, n=64/712) and DENV3 (0.1%, n=1/712) were also detected. Since 2014 a notable increase in DENV2 cases is observed, with frequencies increasing rapidly from 1.03% (n=12/1161) in 2014, to 1.39% (n=23/1650) in 2015, 12.53% (n=53/423) in 2016, 30.56% (n=52/423) in 2017, 71.5 (n=271/379) in 2018 to 87.5% (n=1539/1758) in the first semester of 2019 (**Figure 1A**).

**Figure 1.**
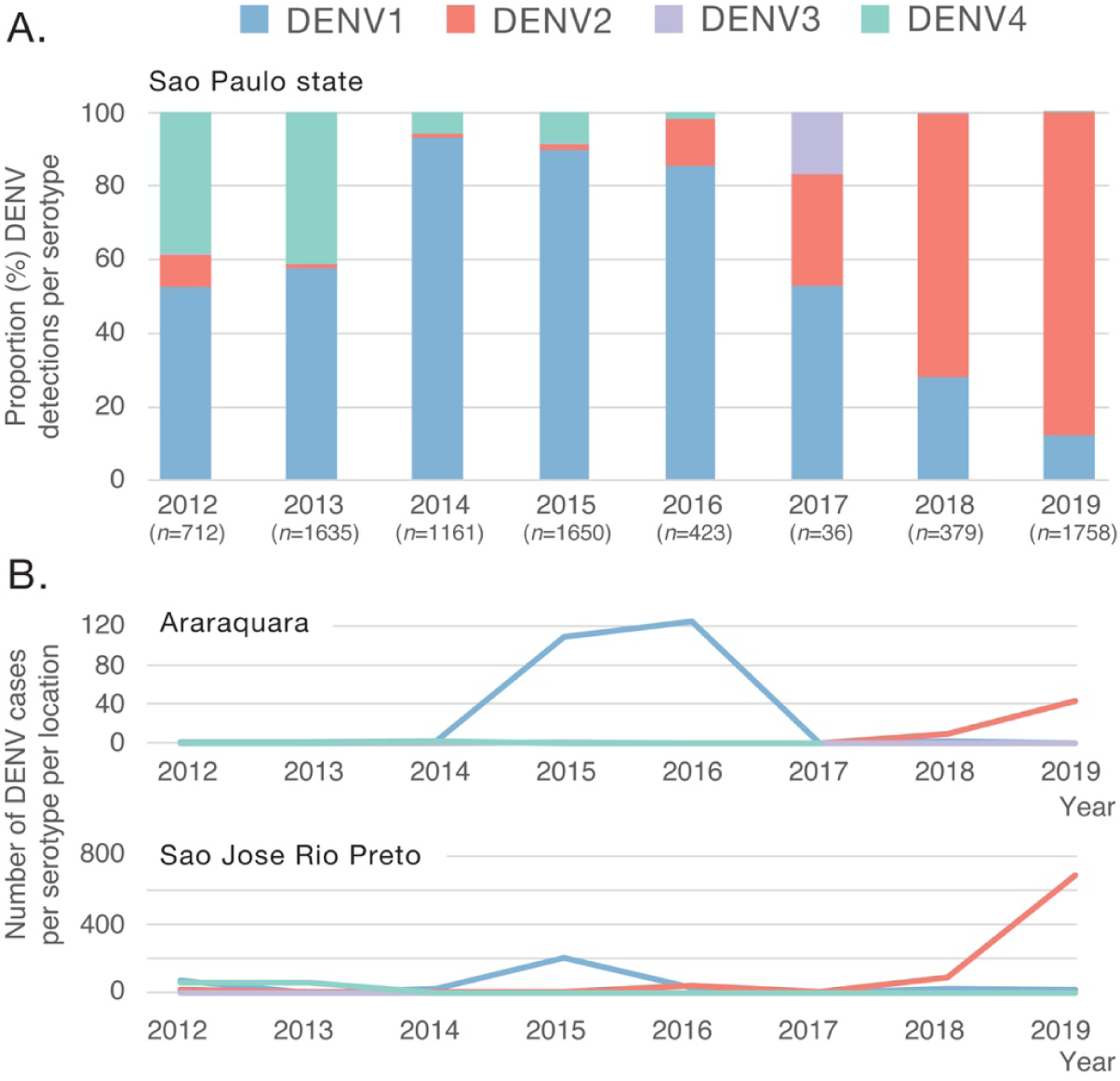
Annual number of DENV cases by serotype reported to the Centro Vigilância Epidemiológica (CVE), Sao Paulo state, Brazil, between January 2012 and June 2019. A. Proportion (percentage) of dengue cases by serotype reported in Sao Paulo state (total *n*=7754). B. Number of dengue cases by serotype in Araraquara (total *n*=294) and in São José do Rio Preto (total *n*=1347).

We next report genomic epidemiological findings from our surveillance of two municipalities in the São Paulo state, Araraquara (ARA) and São José do Rio Preto (SJRP), between early June 2017 and the end of April 2019. In Araraquara, a total of 96% (n=52/54) dengue cases notified to CVE were caused by DENV2 between 2017 and June 2019; in São José do Rio, Preto, during the same period, 95% (n=779/821) were caused by this serotype (**Figure 1B**). From each location, ten RT-qPCR positive samples (mean cycle threshold: 19.8, range: 16.4 to 25) were randomly selected for complete genome sequencing using a previously developed amplicon-based approach for on-site sequencing using the minION platform [24, 25].

All sequenced samples were classified as DENV2 using an automated phylogenetic-based serotyping tool [3]. To investigate the genotypic diversity and the origins of the ongoing DENV2 outbreak in Brazil, we performed phylogenetic analysis of DENV2 using all publicly available complete or partial DENV2 genome sequences (n= 1,630 as of 21 June 2019).

DENV2 is classified into six genotypes, named I-VI [3]. To date, 3 genetic lineages of DENV2 genotype III (DENV2-III) have been reported in Brazil on the basis of phylogenetic analysis of the relationships of partial and complete genomes of circulating strains. These three circulating lineages have been named as lineages 1-3 [5] or BR1-BR3 [26]. Our maximum likelihood (ML) phylogenetic analysis shows a separate introduction of DENV2, hereafter named DENV2-III BR-4 (**Figure 2A**).

**Figure 2.**
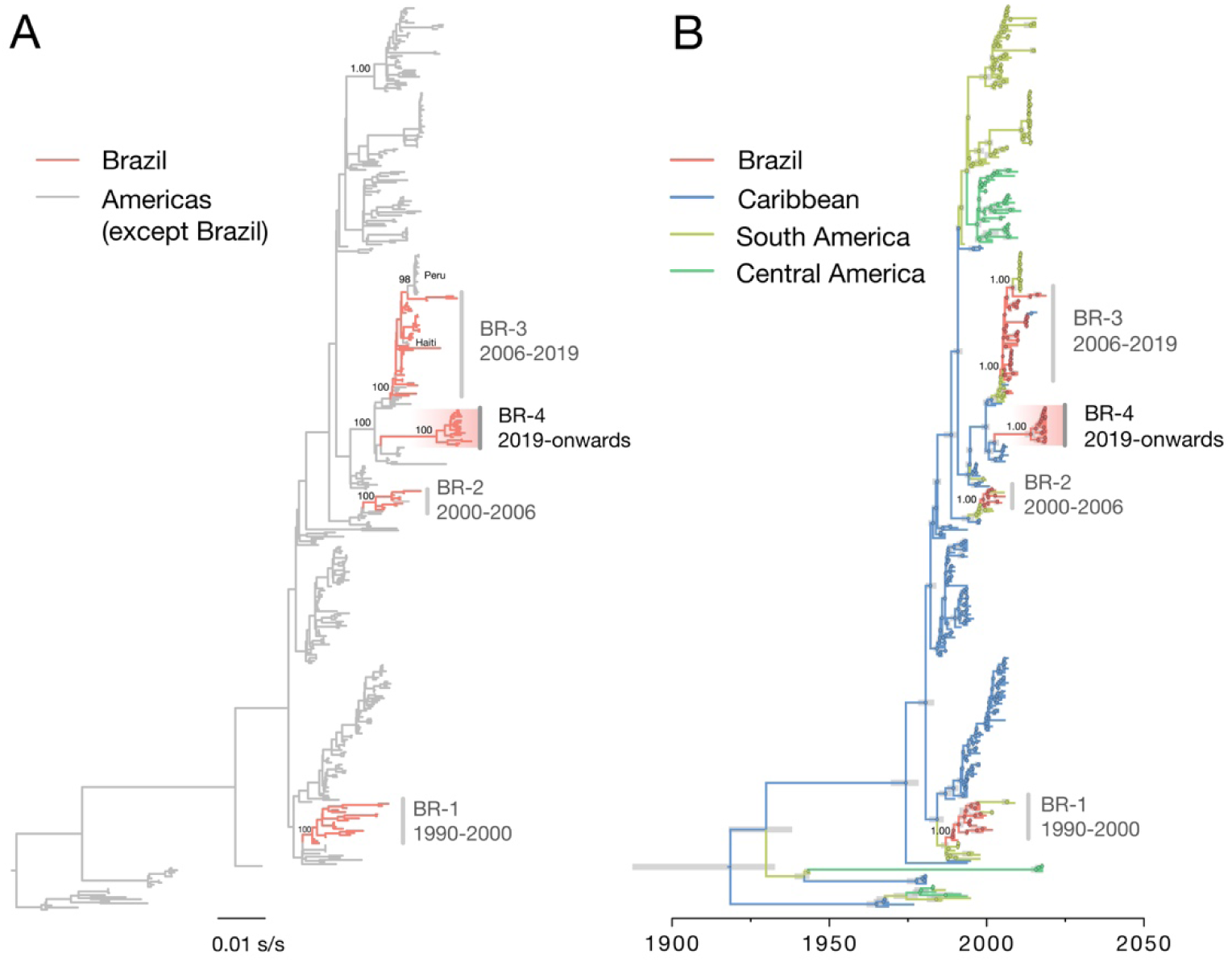
Evolutionary history of DENV2 in Brazil. Maximum likelihood phylogeny (A) and dated phylogeographic tree (B) of DENV serotype 2 (*n*=436) in the Americas. Tips (and nodes leading to) Brazilian strains are shown in red. New clade comprising isolates from 2019 collected in Araraquara and São José Rio Preto (São Paulo state) are highlighted with a red gradient. In panel B, 95% Bayesian credible intervals for node ages are shown for nodes with posterior support above 0.95.

Our ML phylogenetic analysis further shows that virus genomes recovered from the 2019 cases all belong to genotype III (also known as American/Asian genotype). Notably, our analysis strongly supports (approximate likelihood ratio test = 1.00) the clustering of the 2019 DENV2 cases from Brazil (18 of the 19 sequences collected in 2019) into a single monophyletic group (named here as DENV2-III BR-4), which is the result of a new and recent introduction of DENV2-III from outside of Brazil (**Figure 2A**). In addition, two sequences from São José do Rio Preto, one sampled in June 2017 (ID: 126) and another in January 2019 (ID: 140) grouped with the isolates from a previous clade circulating since 2006 in the Southeast Region (BR-3).

To investigate in more detail the origins of the new DENV2-III BR-4 circulating in Brazil in 2019, we conducted a statistical phylogeographic analysis of 436 DENV2 genome sequences representing DENV2 genotypes circulating in the Americas. Our analysis reveals that the DENV2-III BR-4 lineage was introduced in or around 2014 (95% Bayesian credible interval: 2012 to 2015) (**Figure 2B**). From then onwards, the proportion of DENV2 cases in São Paulo state increased (**Figure 1A**). Estimation of the ancestral location of this lineage reveals that it likely originated in the Caribbean region (posterior support = 1.00). However, we note that the new lineage could have also been introduced from countries with tropical climates in South America (e.g. French Guiana or Suriname) from where no recent virus genomic data has been made available. Molecular clock analyses have shown that novel DENV lineages have been introduced in Brazil every 7 to 10 years, after which they are replaced by a novel lineage introduced from other locations. For example, DENV2-III BR-1 was introduced in 1990, DENV2-III BR-2 in 1998, DENV2-III BR-3 in 2005 [5]. Given that the former lineage replacement in Brazil resulted from a strain introduced around 2005, our dating estimate for the introduction of DENV2-III BR-4 and the noticeable increase in number of DENV2 cases in Sao Paulo state are indicative of a new DENV2 lineage replacement in Brazil.

Our data reveals that two distinct virus lineages (DENV2-III BR-3 and DENV2-III BR-4) are co-circulating in a single location (São José do Rio Preto) (**Figure 2**). Although we are limited by the small sample size of the data analysed here, it is remarkable that the frequency of DENV2-III 2019 strains belonging to BR-3 is 5.3% (n=1/19) and for BR-4 is 94.7% (n=18/19). The upsurge in the number of dengue and dengue severe cases observed in São Paulo state, combined with the simultaneous detection of these two lineages and the increased frequency of BR-4 suggests that we are capturing a lineage replacement in real-time.

The surveillance lag between the detection and estimated date of introduction of the DENV2-III BR-4 could result from a lack of genomic surveillance of dengue in a period when Brazil was hit by explosive epidemics of Zika, chikungunya and yellow fever viruses [27, 28]. This highlights the need of improved longitudinal surveillance using, for instance, sequence-independent approaches for arbovirus detection. Such approaches will be particularly critical in geographic regions with year-round DENV transmission in *Aedes* spp. mosquitoes, such as locations with tropical and subtropical climates [2]. Integration of routine serological, molecular, and genomic data along with improved digital surveillance [29], will help to anticipate the arrival and establishment of new virus lineages and other pathogens in the Americas.

## Data availability

XML files and datasets analysed in this study are available in the GitHub repository (https://github.com/arbospread/DENV2). New sequences have been deposited in GenBank under accession numbers XXXXX.

## Acknowledgments

The research was supported by the FAPESP-MRC grant (CADDE, FAPESP 2018/14389-0), a Wellcome Trust and Royal Society Sir Henry Dale Fellowship (grant 204311/Z/16/Z), and by a CNPq # 400354/2016-0 and FAPESP # 2016/01735-2, and by the Oxford Martin School. This work was supported by Decit/SCTIE/MoH and CNPq (440685/2016-8 and 440856/2016-7); by CAPES (88887.130716/2016-00, 88881.130825/2016-00 and 88887.130823/2016-00); by EU’s Horizon 2020 Programme through ZIKAlliance (PRES-005-FEX-17-4-2-33). MLN is supported by FAPESP (Grant # 13/21719-3). ACBT and KRD receive FAPESP Scholarships (Grants #15/12295-0 and 15/14313-6, respectively). MLN and ECS are CNPq Research Fellows (PQ).

